# Dorsal ocelli set the luminance-dependent operating state of the bumblebee visual system

**DOI:** 10.64898/2026.09.15.751776

**Authors:** C. E. Restrepo, Priscila Araujo, Sanja Mikulovic, Pavol Bauer, Emily Baird

## Abstract

Despite long-standing hypotheses that insect dorsal ocelli modulate compound-eye processing to support flight stabilization, dim-light navigation, and locomotor speed, the neurophysiological basis of these functions remains unclear. Combining behavioural assays with multi-site local field potential recordings in bumblebees, we tested how ocellar input affects compound-eye processing. Ocellar occlusion impaired orientation precision at dusk, supporting a role for ocelli in dim-light navigation. Under bright daytime skies, occlusion did not affect orientation but reduced flight speed, consistent with a role in locomotor control under high illumination. Neurophysiologically, ocellar occlusion disrupted luminance-dependent scaling across the visual system, most prominently in the medulla. In intact bees, broadband neural power scaled inversely with luminance, decreasing under bright and increasing under dim conditions. When ocellar input was blocked, this relationship reversed, leaving visual-system activity in a high-power, dark-like state even under bright illumination. Ocellar modulation was particularly evident in the green-sensitive pathway, implicated in optic-flow processing and flight-speed regulation, providing a neural correlate of the behavioural speed reduction while UV-sensitive responses remained largely invariant following ocellar occlusion. These findings reconcile disparate views of ocellar function and identify ocelli as regulators of the neural dynamic range supporting orientation in low light and movement control in bright conditions.

## Introduction

The adult visual systems of most insect orders comprise two categories of eyes–a pair of compound eyes that gather and process the vast majority of visual information and smaller camera-type eyes typically known as the ocelli. Ocelli exist in the most basal insect lineages and likely predate the origin of insects (1). While there is broad intra- and inter-specific variation in their size, number and location, most studied ocellar systems typically provide wide-field sampling of the sky and horizon that is best suited to encoding information about global luminance levels, allowing for rapid head stabilization in flight (2, 3). Another common feature of ocelli is that they are dichromatic with peak sensitivity in the UV and green ranges, an adaptation that has been proposed to support colour constancy (4) and to aid in head stabilization by boosting sky-ground contrast (3). Studies in bumblebees also provide strong evidence that UV-sensitive photoreceptors in the ocelli detect polarized light, a capability that has been hypothesised to underly celestial cue-based orientation behaviour in dim light when compound eye performance deteriorates (5,6). Ocelli have also been implicated in the context-dependent modulation of movement. When moving towards a light source, locusts with occluded ocelli walk slower and follow more tortuous paths than controls, and this effect becomes stronger as light intensity increases (7,8). A cross-species analysis further showed that, in most species, the effect of ocellar occlusion on movement becomes stronger as light intensity increases, with brighter conditions producing larger reductions in speed (9).

Although many studies indicate that information from the ocelli complements that provided by the compound eyes, the neurophysiological basis of how inputs from these two visual systems interact remains unclear. In the context of speed regulation, recordings from the dragonfly ocellar system (10,11) have been interpreted as indicating interactions between ocellar signals, mechanoreceptive input, and signals originating from the compound eyes. However, those studies did not record directly from compound eye pathways, and the influence of compound eye input on ocellar neurons was therefore inferred rather than demonstrated. In the blowfly, motion-sensitive interneurons in the lobula plate have been shown to respond to optic flow while also being modulated by ocellar input (12), providing evidence that information from these two visual systems converges within motion-processing circuits. In locusts, Simmons (13) reported that the descending contralateral movement detector responds to sudden decreases in ocellar illumination, not as an independent light signal, but as a transient facilitatory input that boosts responses to compound-eye visual stimulation. In honeybees, Milde (14) described protocerebral interneurons with more complex response profiles, including antagonistic interactions between ocellar and compound eye input, although the behavioural significance of these responses has not been resolved. In addition, anatomical work in honeybees has reported direct connections between the ocelli and major visual neuropils, including the medulla and lobula, and these pathways have been proposed to contribute to functions such as colour constancy (4). Taken together, these findings suggest that ocellar input can influence particular neuronal circuits within visual pathways receiving compound eye input. However, how ocellar input functionally shapes distributed patterns of population activity across visual brain regions, modulates compound eye processing at a broader network level, and affects behaviour remains unresolved. A major obstacle has been the difficulty of linking behavioural effects directly to circuit-level physiology To address this gap, we combined behavioural assays with electrophysiology to test how ocellar input shapes flight control and compound-eye processing in the bumblebee *Bombus terrestris*. Bumblebees provide a particularly informative model for this question because they navigate across a broad range of light intensities, from the photon-limited conditions of dawn and dusk to the high-intensity midday sun (15,16). This natural behavioural range offers an ideal context in which to test the luminance-dependent transformations long proposed as core functions of the ocelli.

## Results

### Ocellar Contribution to Flight Behaviour Under Natural Sky Conditions

To determine how ocellar input guides flight behaviour across different natural light and celestial regimes, we compared the homing flights (i.e., flights from a feeder to the hive) of control and ocelli occluded bumblebees under clear skies during two recording periods– *daytime* (bright light; sun elevations 32-59°, mean light intensity = 40016 lx) and at *dusk* (dim light; sun elevations 3-11°, mean light intensity = 2371 lx) (Fig. 1A, B). The effect of ocellar occlusion on orientation precision, as indicated by mean vector length, depended on the recording period. Under bright *daytime* conditions, control and ocelli occluded bees oriented with equal precision (F₁ = 1.5, p = 0.2). Yet, at *dusk*, ocelli occluded bees suffered a sharp decline in orientation precision, with significantly shorter mean vector lengths compared to both the ocelli occluded *daytime* flights (χ²₁ = 8.6, p = 0.003) and control flights at *dusk* (χ²₁ = 3.9, p = 0.04). Control bees maintained stable orientation precision across recording periods (χ²₁ = 1.9, p = 0.15), confirming that the performance drop of ocelli occluded bees at *dusk* was directly associated with the loss of ocellar input; (Fig. 2A).

**Fig. 1.**
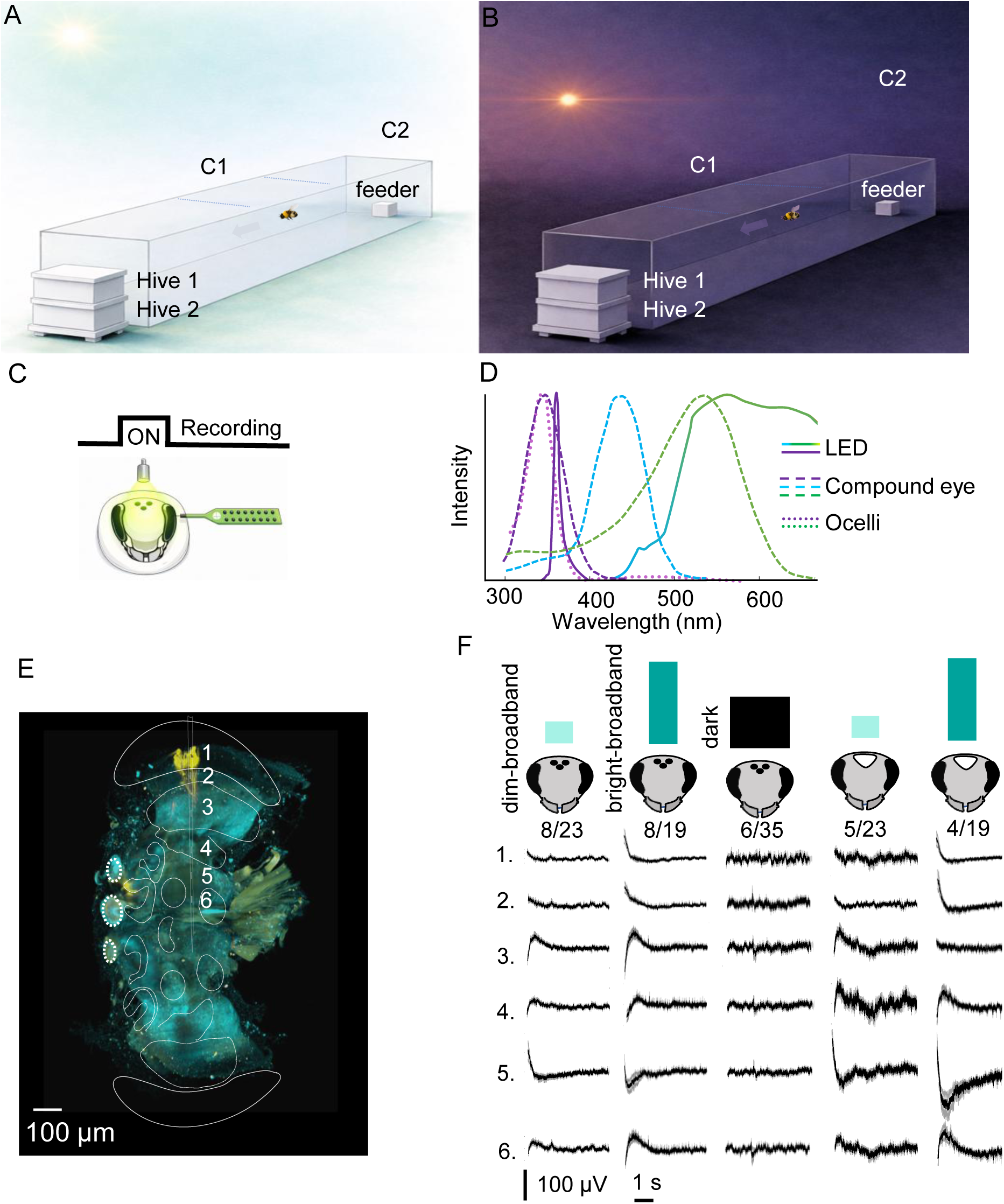
Experimental paradigm for behavioral analyses of ocellar function in bumblebees. (A and B) Outdoor flight arena used for behavioral experiments under *daytime* (A) and *dusk* (B) conditions. Bumblebees were trained to fly between their hives and a feeder at the opposite end of the arena. Flights from the feeder to the hive (*i.e.*, homing flights) were recorded by two overhead cameras (C1 and C2) and one camera near the feeder. (C) Experimental methods for electrophysiological analyses of ocellar function in bumblebees. Electrophysiological preparation for local field potential (LFP) recordings. A 16-channel linear silicon probe was inserted laterally through the eye, and visual stimulation consisted of a 5 s light ON period followed by a 5 s OFF period. All electrophysiological analyses reported in this study were performed on recordings obtained during the OFF period. (D) Spectral composition of the visual stimuli relative to the photoreceptor classes in the bumblebee compound eye (dashed lines) and ocelli (dotted lines) (6, 30). Stimuli were generated using a narrowband UV LED and a white LED with a dominant green component and minor blue output (solid lines). (E) Histological reconstruction of the probe trajectory across the visual pathway. Six recording sites were selected for analysis: retina (1), lamina (2), medulla (3), lobula (4), and two protocerebral sites (5&6). (F) OFF LFP traces recorded simultaneously across the six analysed sites under the main stimulation conditions. Dark mean and gray SEM. Fractions correspond to animals/trials per condition.

**Fig. 2.**
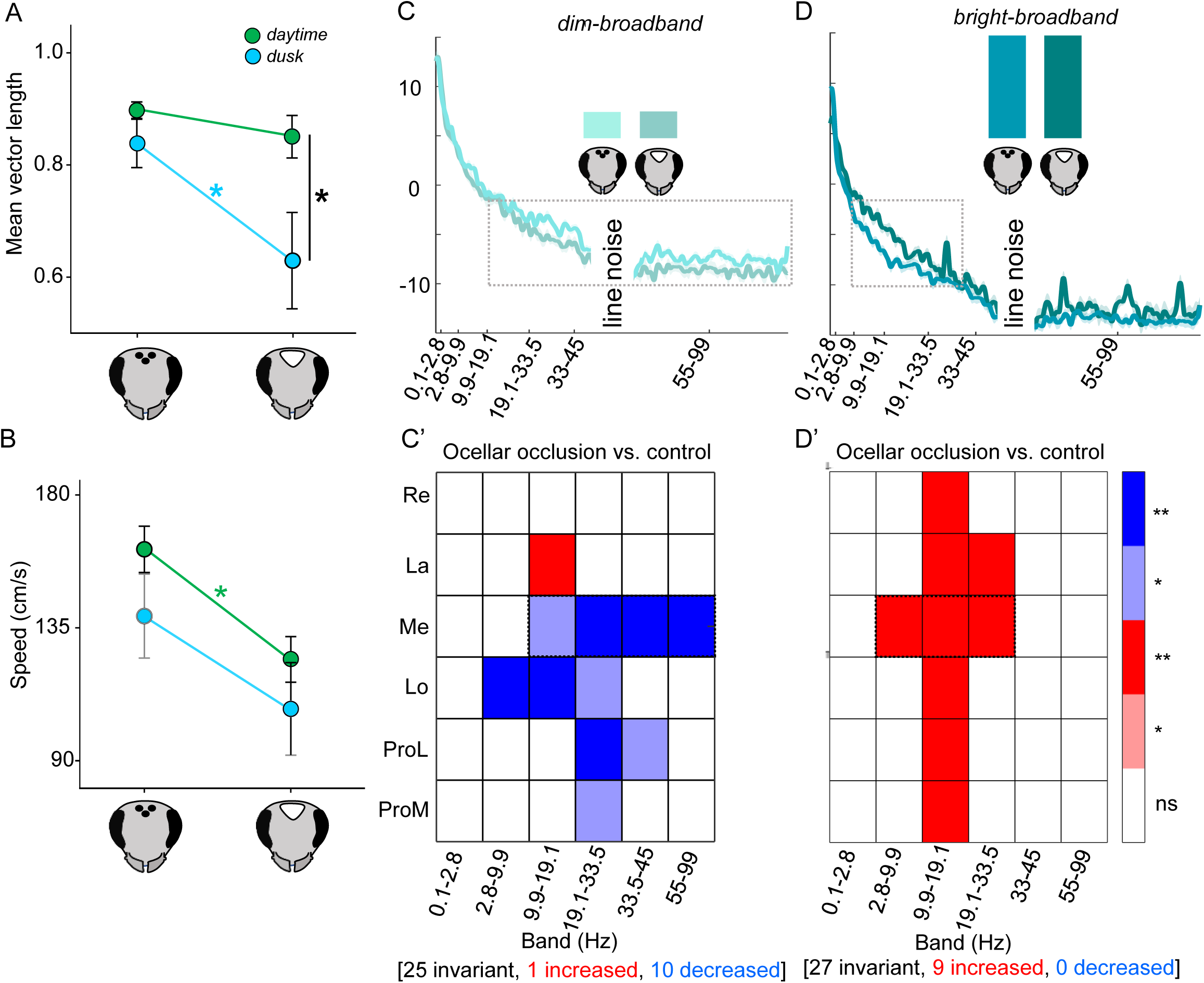
Ocellar occlusion produces complementary light-dependent effects on flight behaviour and visual-system activity. (A, B) Effects of ocellar occlusion on homing-flight behaviour under natural daytime and dusk conditions. (A) Orientation precision, quantified as mean vector length, was unaffected by ocellar occlusion during daytime flights (F₁ = 1.5, *p* = 0.2), but was significantly reduced at dusk compared with control flights (χ²₁ = 3.9, *p* = 0.04). Orientation precision in ocelli-occluded bees was also lower at dusk than during daytime (χ²₁ = 8.6, *p* = 0.003), whereas control bees maintained similar orientation precision across the two light regimes (χ²₁ = 1.9, *p* = 0.15). (B) Flight speed showed the complementary pattern: ocellar occlusion significantly reduced flight speed under daytime conditions (F₁ = 11, *p* = 0.001), whereas control and ocelli-occluded bees flew at similar speeds at dusk (χ²₁ = 0.7, *p* = 0.39). (C, D) Medulla power spectral density (PSD) under *dim-broadband* (C) and *bright-broadband* (D) stimulation in control and ocelli-occluded bees. Under dim-broadband stimulation, ocellar occlusion reduced medulla PSD, particularly across approximately 9.9–99 Hz. Under bright-broadband stimulation, the direction of the effect was reversed: ocellar occlusion increased PSD relative to controls, most prominently across the low- to mid-frequency range. The 45–55 Hz interval was excluded because of line noise. (C′, D′) Heat maps summarize the effect of ocellar occlusion across six brain regions and six frequency bands. Values represent the difference between control and occluded conditions (ΔPSD = control − occluded). Blue indicates frequency bands in which PSD was lower after ocellar occlusion, red indicates bands in which PSD was higher after occlusion, and white indicates no detectable difference. Under *dim-broadband* stimulation (C′), occlusion predominantly reduced PSD [25 invariant, 1 increased, 10 decreased], with the strongest and broadest effects centered in the medulla and weaker effects in the lobula and protocerebrum. Under *bright-broadband* stimulation (D′), occlusion instead increased PSD [27 invariant, 9 increased, 0 decreased], with effects centered in the medulla but extending into the retina and lamina, particularly across approximately 2.8–33.5 Hz. Light shading denotes *p* < 0.10, dark shading denotes *p* < 0.05, and white denotes nonsignificant comparisons. Re, retina; La, lamina; Me, medulla; Lo, lobula; ProL and ProM, lateral and medial protocerebral recording sites.

Interestingly, flight speed exhibited the exact opposite dependency: ocelli occluded bees flew slower than controls but only under *daytime* conditions (F₁ = 11, p = 0.001; Fig. 2B). At *dusk*, both control and ocelli occluded bees flew at similar speeds (χ²₁ = 0.7, p = 0.39). Within treatments, there was no strongly significant effect of recording period on flight speed (control: F₁ = 2.2, p = 0.13; ocelli occluded: χ²₁ = 3.8, p = 0.05), although there was a tendency for flights at *dusk* to be slower than those under *daytime* conditions.

To uncover the circuit-level basis of these behavioral shifts, we performed neurophysiological recordings using a broadband stimulus (i.e., one that contained peaks in the UV and green, overlapping with the sensitivity profiles of ocellar photoreceptors (6), Fig. 1C, 1D) on control and ocelli occluded bees at different light intensities. Specifically, we tested whether ocellar inputs generate distinct neural signatures in the compound eye processing pathways that could account for the impact that ocellar occlusion had on flight behavior under *daytime* and *dusk* conditions.

### Ocellar dependent gain control and luminance normalization in the medulla

We first verified that the *dim-broadband* stimulus reliably activated the visual pathway by comparing its power spectral density (PSD) to the *dark* baseline (Figure 2—figure supplement 1). These PSD comparisons across frequency bands and visual regions led to 12 invariant, 12 increased and 12 decreased responses, confirming that this intensity was sufficient to evoke neural responses (hereafter, the outcome of band/region PSD comparisons will be presented as [# invariant, # increased, # decreased]). We next compared the *bright-broadband* stimulation to darkness (Figure 2—figure supplement 1B) and, once again, found thresholded differences but, in comparison to the *dim-broadband/dark* comparison, the overall trial-level PSD tended to decrease [15, 7, 14]. These results indicate that both our *dim-broadband* and *bright-broadband* stimuli elicited robust and distinct post-stimulus (OFF) patterns relative to the *dark* baseline, confirming that the effects we observed reflect stimulus-evoked modulation rather than time-dependent baseline fluctuations in darkness.

To investigate how the ocelli modulate compound eye circuits, we recorded LFPs in control bees and then re-recorded from the same individuals after ocellar occlusion (denoted hereafter by *-_OC_)*, comparing intact and ocelli occluded responses under low- and high-luminance conditions. Although the laboratory stimuli were not photon-equivalent to the luminance during the *dusk* and *daytime* periods in the behavioural experiments, they provided a physiological analogue of the contrast between these conditions: Light intensity under *dusk* conditions was 95% lower than under daytime conditions and the *dim-broadband* stimulus was 98% lower than the *bright-broadband* stimulus. Under *dim-broadband* stimulation, ocellar occlusion produced an overall decrease in PSD [25, 1, 10] (Fig. 2C) with the pattern of thresholded band/region effects concentrated in the medulla and weaker impacts in the lobula and protocerebrum (Fig. 2C’). Under *bright-broadband* stimulation (Fig. 2D), ocellar occlusion instead produced an overall increase in PSD [27, 9, 0], with prominent effects extending into the retina and lamina while remaining centred on the medulla (Fig. 2D’).

We next compared the effects of luminance, that is, *dim-* versus *bright-broadband* stimulation within each treatment group (Fig. 3). In control bees (Fig. 3A), the most consistent effect was a suppression of power across low-, mid-, and high-frequency ranges [20, 0, 16], especially within the compound eye neuropils (retina, lamina, and medulla). The protocerebrum showed only a limited suppression at mid-range frequencies, while the lobula appeared largely invariant. When the ocelli were occluded, this pattern was markedly altered (Fig. 3B). Global brain activity showed reliable modulation [23, 9, 4], again centered around the medulla, with almost full invariance in the retina and lamina. Taken together, these trial-level comparisons revealed a consistent pattern of modulation across region–band combinations, with effects accumulating most strongly in the medulla. We therefore asked whether this apparent anatomical concentration was supported by enrichment analysis. Structure-level enrichment confirmed the dominant effect to the medulla (Fig. 4A), and medulla-restricted enrichment showed that this modulation was concentrated primarily in the 2.8–33.5 Hz range (Fig. 4B).

**Figure 3.**
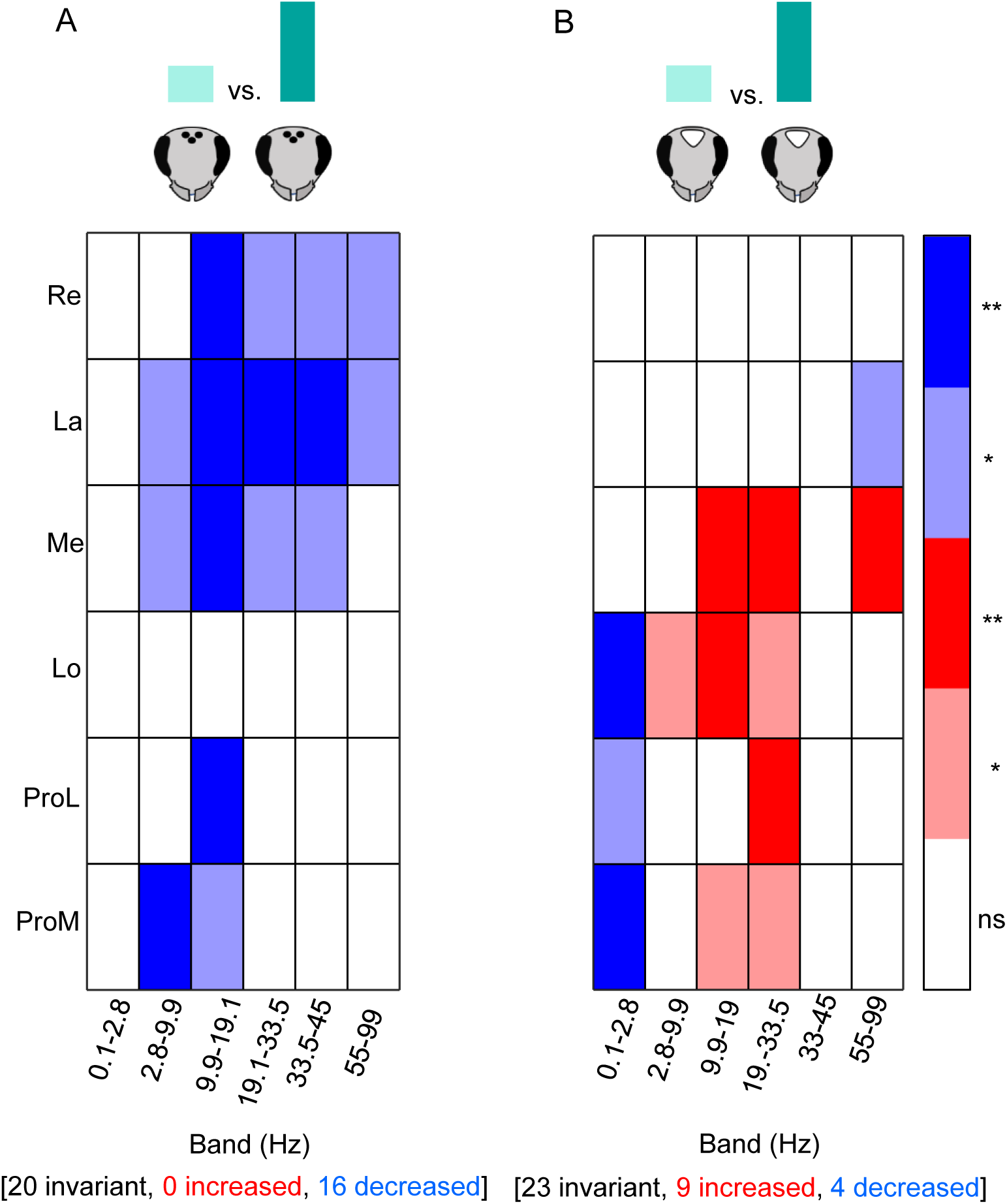
Ocellar occlusion disrupts luminance-dependent suppression of visual brain activity. (A)Intact ocelli. Increasing light intensity produced broad suppression of low- and mid-frequency power (2.8–45 Hz) across retina, lamina, and medulla, with additional suppression in the protocerebrum. Very low frequencies (0.1–2.8 Hz) remained largely unchanged. This pattern reflects a coordinated gain-control mechanism that downscales neural activity as luminance rises. (B) Ocelli occluded. When the ocelli were blocked, this suppressive pattern disappeared. Instead, medulla and lobula channels showed widespread enhancements at mid–high frequencies (9–99 Hz) although stronger in the medulla and only isolated sites displayed weak low-frequency suppression. Noted also that the retina and lamina activity is practically the same when the ocellies were occluded.

**Fig. 4.**
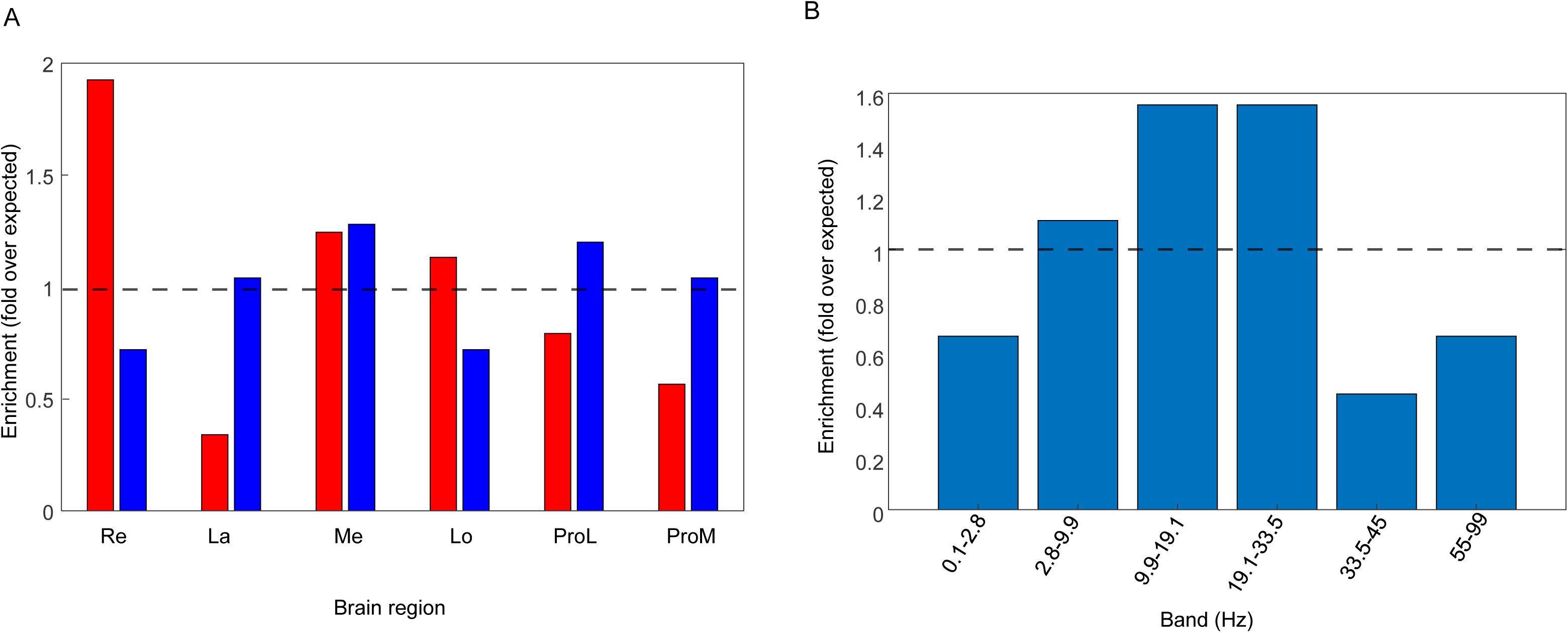
Enrichment analysis identifies the medulla and its mid-frequency bands as the dominant locus of modulation. (A) Structure-level enrichment index for thresholded PSD modulation across the visual pathway, increases (red) and decreases (blue). Values are normalized to the brain-wide fraction of thresholded region × band effects, such that 1 indicates the expected background level of modulation. The medulla shows the highest enrichment for both increases and decreases in PSD, identifying it as the principal hotspot of luminance-dependent modulation. The retina also exhibits elevated enrichment driven by increases, while the lamina and central protocerebrum remain strongly reduced, reflecting their largely invariant responses across illumination conditions. (B). Frequency-specific enrichment of modulation in the medulla. Bars show the enrichment index for each frequency band within the medulla, defined as the band-specific fraction of thresholded changes normalized to the medulla-wide average (dashed line = 1, expected level). Mid-frequency bands (9.9–19.1 Hz and 19.1–33.5 Hz) were strongly over-represented, whereas lower and higher frequencies were comparatively stable. This indicates that medulla dominance arises primarily from modulation in the 2.8–33.5 Hz range.

Having established the medulla as the main site of this effect, we next examined how the medulla PSD was organized across luminance conditions relative to a common reference state. To do this, we normalized all medulla conditions to the *dark* baseline, allowing us to determine whether each stimulus increased or decreased band-limited power relative to the same reference and, crucially, whether ocellar input altered this relationship. This analysis revealed a clear luminance-dependent ordering of medulla activity (Fig. 5). With ocelli intact, PSD scaled inversely with luminance, such that *dark* was associated with higher power, *dim-broadband* stimulation with intermediate values, and *bright-broadband* stimulation with the lowest values. In contrast, when ocellar input was removed, this organization collapsed and the medulla PSD ordering inverted; *bright-broadband* stimulation no longer occupied the low-power end of the response range but instead shifted the medulla toward *dim-broadband* and *dark* values.

**Fig. 5.**
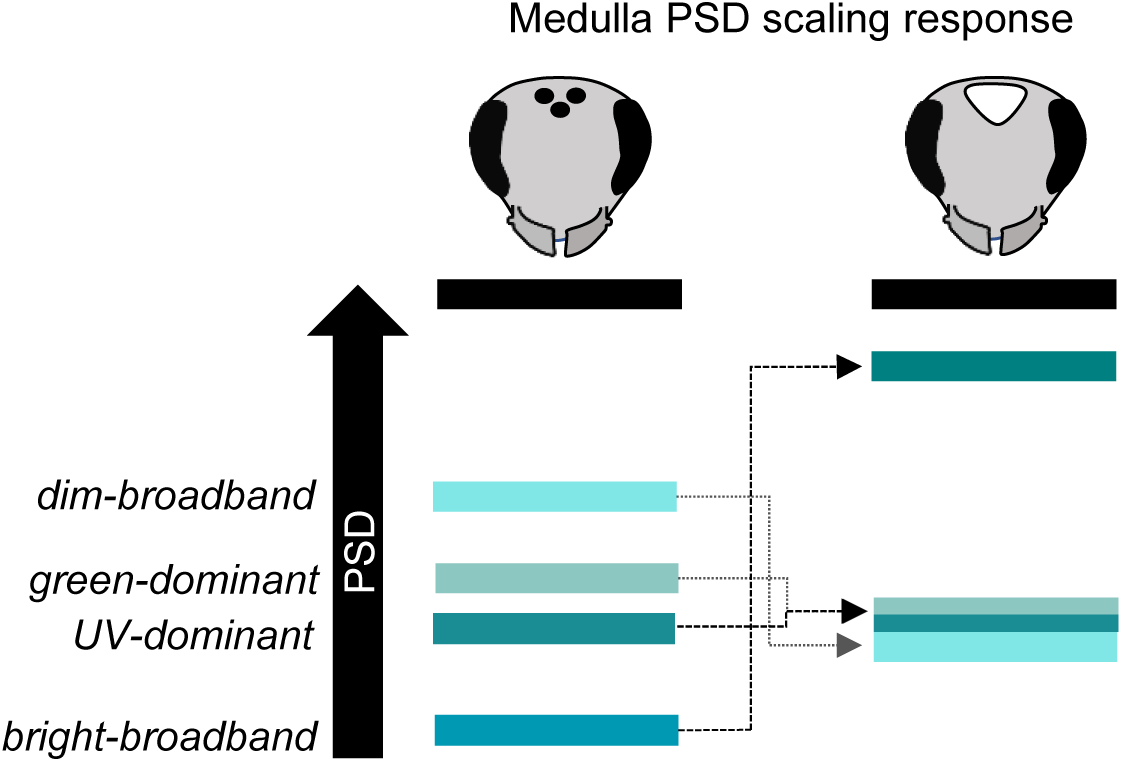
Ocellar occlusion disrupts the normal luminance-dependent ordering of medulla activity. Medulla activity (channel 7) was quantified as the mean PSD in the 2.8–33.5 Hz band, normalized within each trial by the mean PSD across the analyzed spectrum and then expressed relative to the Dark condition (Dark = 1). In intact animals (left), the medulla showed a normalization-like scaling pattern, with relative band weighting progressively decreasing from Dark toward *dim-broadband* and then further toward the mixed spectral states, reaching its lowest level under *bright-broadband* stimulation. After ocellar occlusion (right), this ordered scaling was reorganized: *bright-broadband_oc* shifted upward toward dark-like, whereas *dim-broadband_oc* and the mixed spectral states remained in a lower, compressed range.

### Synaptic and network contributions dominate luminance-dependent medulla activity

Given that our enrichment analysis identified the medulla as the structure most strongly modulated across illumination conditions, we next asked how much of its luminance-dependent activity could be accounted for by neuronal output. Specifically, we tested whether medulla LFP power covaried with detected spikes or with multi-unit activity (MUA), reflecting the combined activity of local neuronal populations (Figure 5—figure supplement 1). Detected spikes showed only weak associations with band-limited LFP power under all conditions, with mean ρ² values ranging from 0.0005 to 0.0162, indicating that threshold-detected spike activity did not closely track the observed LFP dynamics. By contrast, MUA was more frequent in the dark and became more tightly coupled to LFP power under dim and bright illumination, consistent with stronger synchronization between local population firing and field activity. MUA showed substantially stronger associations with band-limited LFP activity than detected spikes, with mean ρ² values ranging from 0.1695 to 0.3479 across conditions (16.95–34.79% when expressed as ρ² × 100; Figure 5—figure supplement 1C), consistent with observations in Drosophila (18). Even so, these associations remained moderate, indicating that much of the band-limited LFP dynamics were not closely associated with local population firing. Together, these results indicate that the luminance-dependent LFP changes in the medulla are not driven primarily by spike output, but instead likely reflect dominant synaptic and broader network processes. This pattern is consistent with observations in the primate visual cortex (17), where LFP power and MUA are moderately but not fully correlated, suggesting that, in our experiments, the OFF-LFP activity captures broader synaptic processes.

### Ocellar modulation of early compound-eye pathways involved in optic flow

In ocelli-occluded bees, activity in the early visual neuropils–the retina and lamina–showed little luminance-dependent modulation. When *dim-broadband-_OC_* and *bright-broadband-_OC_* stimulation were compared (Fig. 3B), most channel-band combinations were invariant, with only a small residual effect in the lamina at 55-99 Hz. This indicates that, without ocellar input, the retina and lamina largely lose their normal differential PSD response to brightness. Behaviourally, this flattening is consistent with the *dusk* condition, where ocelli-occluded bees flew at the same speed as controls (Fig. 2B), suggesting that under low luminance the absence of ocellar input does not further reduce an already slow flight state. A similar pattern emerged when controls were compared with ocelli-occluded bees under *dim-broadband* stimulation (Fig. 2C’): the retina and lamina again showed mostly invariant responses. Although the retina and lamina were largely insensitive to luminance in ocelli occluded bees, comparing control and ocelli occluded bees under *bright-broadband* stimulation revealed clear changes in these same early visual neuropils (Fig. 2D’). Importantly, we found that ocellar modulation produced its strongest effects with the *green-dominant* stimulus, since the pair *green-dominant-_OC_* vs *green-dominant* resulted in a broad activity across the brain, although relatively less in the medulla [20, 0, 16], while its counterpart *UV-dominant-_OC_* versus *UV-dominant* comparison was entirely invariant [36, 0, 0] (Fig. 6).

**Fig. 6.**
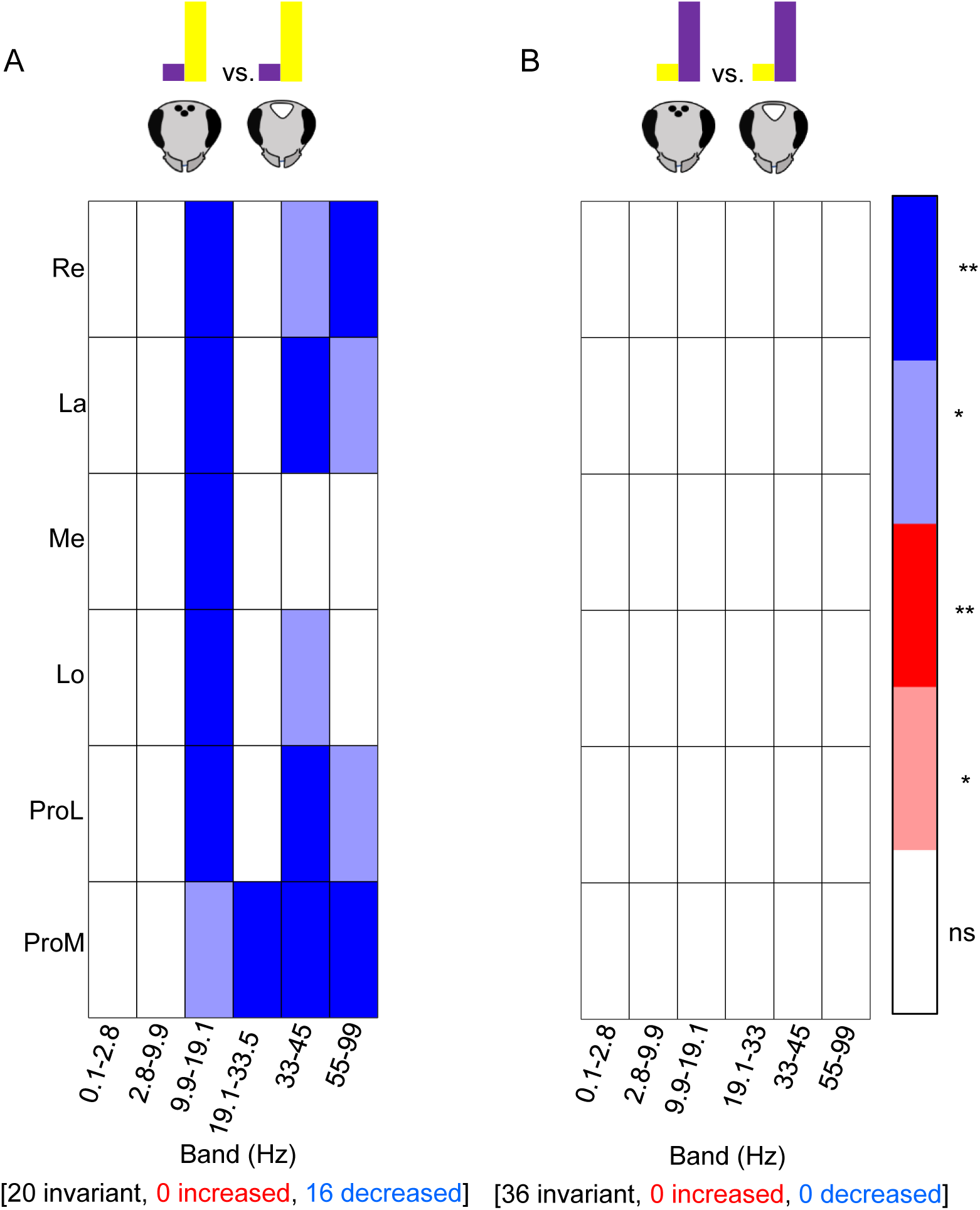
Spectrally biased ocellar effects differ between green-dominant and UV-dominant stimulation. (A) Comparison of green-dominant versus green-dominant-oc conditions. (B) Comparison of UV-dominant versus UV-dominant-oc conditions. Ocellar occlusion produced broader effects under green-dominant stimulation, particularly in first and second order visual neuropils, whereas the UV-dominant comparison was largely invariant, indicating that ocellar modulation of early visual processing is expressed more strongly through the green-sensitive pathway than through the UV-sensitive pathway.

## Discussion

Our behavioural data suggest a distinct, light intensity-dependent dissociation in the role that ocelli play in guiding movement. During the *daytime*, bumblebees with their ocelli occluded were able to maintain a similar orientation precision to controls, but their flight speed was reduced. At *dusk*, ocellar occlusion caused a decrease in orientation precision with respect to controls but did not affect flight speed. This suggests that, in bright daytime conditions, the ocelli facilitate the processing of rapid luminance cues required for high-speed optic flow processing, whereas in dim dusk conditions, they become critical sensors for maintaining stable orientation. To link these behavioral shifts to underlying network dynamics, we utilized multielectrode LFP recordings to capture population activity across the brain (18) and used enrichment analysis to identify where modulation converged most strongly. In agreement with studies in *Drosophila* (19) and bumblebees (20), we identified the medulla as the primary locus for modulation because significant effects repeatedly accumulated there across multiple luminance- and ocelli-dependent analyses.

A central finding of our study is that ocellar input appears to modulate the medulla by a gain control like mechanism. In control bees PSD scaled inversely with light intensity but, after ocellar occlusion, this relationship was inverted so that bright stimulation evoked a dark-like high-power state (Fig. 5). This physiological signature of ocellar occlusion coincides with the loss of orientation precision observed in ocelli occluded bees flying at *dusk* (Fig.2A) and supports the hypothesis that ocelli provide a global luminance reference that preserves visual guidance when contrast is poor (6). This interpretation is also consistent with work in *Drosophila* showing that turning responses are scaled primarily to contrast rather than to absolute luminance across a broad range of light levels (21). Because dusk conditions combine low luminance with weak visual contrast, directional coding should be especially sensitive to failures in luminance calibration. Our results therefore suggest that ocellar input helps stabilize a dim-appropriate medulla state, thereby preserving orientation when visual contrast is poorest.

We further show that ocellar occlusion selectively reduces flight speed during the *daytime* but not at *dusk* (Fig. 2B). Consistent with this behavioural effect, the retina and lamina of ocelli-occluded bees showed little differential activity between dim- and bright-broadband stimulation (Fig. 3B), indicating that removal of ocellar input largely abolished the normal luminance-dependent modulation of these early compound-eye pathways. Thus, the key result is not simply that retinal and laminar activity remained invariant, but that the distinction between dim- and bright-dependent states was lost when ocellar input was removed. This loss of luminance-dependent differentiation may impair the ability of early motion-processing circuits to adjust their operating state under bright conditions, thereby contributing to the reduction in daytime speed. This interpretation is biologically plausible because bumblebee ocelli contain both UV- and green-sensitive photoreceptors (6), the strongest spectral components of the broadband stimulus used in our experiments (Fig. 1D). In the compound eye, green-sensitive photoreceptors provide the fast achromatic input used for early motion detection (22), with the lamina forming the first stage of this pathway (23), including in bumblebees (24). These findings suggest that the principal ocellar influence on early visual processing is exerted through the green channel. Since our broadband stimulus was dominated by green light, and the laminar pathways in bumblebees are associated with the green channel and optic-flow processing, the observed reduction in flight speed likely reflects ocellar modulation of motion-processing circuits. The fact that the dominant effect of the ocelli is conveyed by green and not UV (Fig. 6) supports this interpretation.

The anatomical source of both forms of modulation remains an open question. Rence et al. (25) reported that ocellar occlusion significantly reduced compound eye ERG amplitude, potentially via a direct afferent projection into the lamina. While such direct connectivity has not yet been confirmed in bumblebees, an alternative explanation could involve the extensive network of gap junctions within the visual neuropils (26). Even if ocellar afferents primarily reach the medulla (4), these electrical networks could facilitate retrograde or lateral propagation of activity toward the lamina. Such circuit interactions would be consistent with the established role of ocelli in speed control and stabilization circuits. However, our finding that ocellar input modulates the earliest stages of the visual pathway—the retina and lamina—suggests a broader functional influence than previously recognized.

The effects of ocellar occlusion observed here are conceptually consistent with recent work in *Drosophila* where early lamina neurons distribute contrast and luminance information in parallel: L2 provides a major fast contrast-sensitive input to motion pathways, whereas luminance-sensitive inputs from L1 and especially L3 contribute gain correction that stabilizes motion-guided behavior as illumination changes. Rapid luminance gain control is then implemented downstream in medulla neurons such as Tm1 and Tm9, which normalize contrast responses before they reach direction-selective circuitry (19). Our results suggest that bumblebees (and possibly other insects) may solve a related problem using an additional source of information: the ocelli. In intact bees, medulla activity scaled inversely with luminance, whereas after ocellar occlusion this organization collapsed and bright stimulation shifted toward dim- and dark-like states. This pattern is consistent with the idea that ocellar signals provide an extrinsic luminance reference that helps stabilize motion-relevant visual processing across changing light conditions. Such a mechanism would be particularly valuable in bumblebees, which fly across a broad luminance range.

Importantly, the underlying circuitry in bees may not be identical to that of flies. Bumblebee lamina cells show physiological properties that differ from the standard fly description (24), suggesting that luminance–motion interactions may be implemented somewhat differently in Hymenoptera. Although previous studies in *Drosophila* did not assess the role of ocelli, the bee lamina is not a passive relay and could plausibly participate in the ocellar modulation inferred from our study.

Several limitations bound the specificity of our mechanistic claims. First, restricting our analysis to the OFF condition limits our inferences regarding stimulus-locked dynamics and early transient components, although Kondo et al. (10) reported that lateral ocellar nerves were inhibited during illumination and discharged in the OFF phase. Second, our interpretation relies mostly on the pattern of PSD change across channel-band combinations, rather than on specific frequency bands, because we did not identify a single unique oscillatory correlation of luminance. Third, our broadband and spectrally biased stimuli likely engaged circuits that jointly support navigation, landmark processing, color constancy, and optomotor stabilization; thus, the observed patterns reflect integrated activity across overlapping functional domains. Although ocellar occlusion clearly impairs directional stability (27), flight stabilization has classically been associated with large second-order ocellar interneurons and downstream descending pathways that convey ocellar signals to thoracic flight-control circuits (28). The medullary modulation we observe therefore likely reflects a parallel route, separate from the canonical ocellar pathway involved in attitude stabilization. Finally, because our electrophysiological comparisons were performed at the trial level, these analyses should be interpreted as exploratory evidence of effect direction, consistency, and anatomical organization rather than as formal population-level inference across animals. Nevertheless, the convergence of effects across luminance manipulations, visual neuropils, spectral conditions, and behavior provides a coherent hypothesis-generating framework for understanding how ocellar input interacts with compound-eye processing.

## Materials and Methods

### Animals and behavioral assays

Bombus terrestris foragers obtained from commercial colonies (Koppert Biological Systems, the Netherlands) were used for all experiments. Behavioral testing was performed in an outdoor flight arena (1.1 m × 3.5 m × 1.1 m). Bees from 2 stacked hives, with entrances at heights of 15 cm and 85 cm, were trained to fly to a feeder containing 50% sucrose solution at the opposite end of the arena. Homing flights were defined as uninterrupted flights from feeder to hive and were recorded during morning, midday, and dusk sessions. For analysis, recording periods were grouped as daytime (sun elevation 32°–59°) and dusk (3°–11°). Only flights recorded under clear skies were included.

To test the effect of ocellar occlusion, baseline control flights were recorded over 3 d. Ocelli were then occluded with black and white matte paint, after which bees were returned to the arena and re-recorded following a 24-h recovery period (Fig. 1A-B)..

Flight trajectories were recorded with 3 synchronized cameras (2 overhead and 1 near the feeder) and digitized in 2 dimensions using DLTdv8a in MATLAB. Checkerboard calibration was used to correct fisheye distortion and convert trajectories to metric coordinates. Within a 1-m central tracking zone, we quantified mean vector length (r) and flight speed (cm/s). Behavioral statistics were performed in R using linear models and linear mixed-effects models, with bee identity included as a random effect where appropriate.

### Electrode preparation and insertion and electrode locations in the brain

Recordings were obtained with a 16-channel linear silicon probe (A1x16-3mm100-177; NeuroNexus) coated with Texas Red dextran. Bees were secured in a tube with dental wax, and the bases of the antennae were immobilized. Using a micromanipulator, the probe was inserted laterally through the eye, perpendicular to eye curvature, along a trajectory crossing the lamina, medulla, and lobula. Signals were acquired with an INTAN RHD USB interface board at 25–30 kHz. Based on body size and histological verification, 6 recording sites were selected for analysis: retina, lamina, medulla, lobula, lateral protocerebrum, and medial protocerebrum (Fig. 1C).

### Visual stimuli

Visual stimuli were delivered with a narrowband ultraviolet (UV) LED and a white LED with a dominant green component and minor blue output (Fig. 1D). Stimuli consisted of 5 s ON followed by 5 s OFF. Four light conditions were used: dim-broadband, bright-broadband, green-dominant, and UV-dominant. A dark condition was recorded after 30 min in darkness. Ocellar-occluded conditions are indicated by the suffix ‘-_OC_’.

Experimental conditions were defined as follows:

- dim-broadband: both LEDs at low intensity, 1.25 × 10^14 photons/s/cm²
- bright-broadband: both LEDs at high intensity, 5.25 × 10^15 photons/s/cm²
- green-dominant: white LED high, UV LED low, 5.0 × 10^15 photons/s/cm²
- UV-dominant: UV LED high, white LED low, 2.82 × 10^15 photons/s/cm²

### Local field potential recording and analysis

Raw extracellular recordings were preprocessed in MATLAB using EEGLAB functions, following the general logic of the multichannel insect-brain pipeline described by Paulk et al. (2013). Intan files were first imported, and channels were reordered to match the physical layout of the silicon probe. To isolate the local field potential (LFP) range, signals were low-pass filtered at 300 Hz in EEGLAB while retained at their native acquisition sampling rate (25 or 30 kHz). Independent component analysis (ICA) was then applied to the filtered multichannel data to reduce shared artifacts and improve separation of neural components before resampling. The ICA-processed signals were subsequently resampled to a common sampling rate of approximately 6.67 kHz and corresponding downsampled time vectors were generated. To identify persistently noisy channels, all down sampled ICA-processed segments belonging to the same recording block were concatenated and screened jointly in EEGLAB using pop_rejchan with a normalized probability criterion (threshold = 2). Channels identified as bad were replaced with NaN, and the cleaned concatenated dataset was then split back into its original trial segments and saved for subsequent LFP spectral and coherence analyses. Spike and MUA analyses were performed separately on the corresponding broadband extracellular recordings, as described below. This procedure preserved the anatomical organization of the recordings while reducing shared noise and maintaining stable channel quality for downstream analyses.

We carried out a clustering analysis using *k*-mean on the LFP power spectra to identify data-driven frequency domains present in endogenous brain activity. K-means clustering identified six data-driven frequency bands, corresponding approximately to 0.1–2.8 Hz, 2.8–9.9 Hz, 9.9–19.1 Hz, 19.1–33.5 Hz, 33–45 Hz, and 55–99 Hz (Figure 1—figure supplement 1). These data-driven frequency bands provide a biologically grounded framework for subsequent analyses of spectral encoding under different light and UV conditions.

### Power spectral density (PSD) analysis and statistics

Power spectral density (PSD) was estimated from 0.025 to 99 Hz using Welch’s method with overlapping windows. Power values were normalized within each trial and converted to decibel (dB) units to facilitate comparison of spectral profiles across recordings and preparations. Data-driven frequency bands were subsequently defined from the global LFP spectrum. Analyses were restricted to the post-stimulus OFF period because the stimulus ON period contained LED-related artifacts, whereas OFF responses showed robust and reproducible post-stimulus activity.

For each brain region × frequency-band combination, differences in band power between experimental conditions were assessed at the trial level using Wilcoxon rank-sum tests. Comparisons were treated as unpaired because individual trials were not matched one-to-one across conditions, including comparisons in which recordings were obtained from the same preparation before and after ocellar occlusion. For each comparison, both the direction of the effect (increase or decrease in power) and its magnitude were recorded. Because multiple trials were collected from the same bee/preparation, trial-level observations were not considered independent biological replicates for formal population-level inference across animals. Instead, these analyses were used to characterize the direction, consistency, and anatomical and frequency-specific distribution of condition-dependent spectral changes across repeated recordings within preparations. Results were summarized as heatmaps of PSD changes across brain regions and frequency bands, with annotations based on raw p-value thresholds (main threshold, *p* < 0.10; stricter threshold, *p* < 0.05).

Enrichment analysis

To determine whether luminance- and ocelli-dependent PSD effects were preferentially concentrated in particular neuropils, we adapted the logic of over-representation analysis to the multiregion LFP dataset. For each neuropil, we calculatedthe fraction of region × frequency-band comparisons showing thresholded modulation and normalized this value by the corresponding brain-wide fraction of thresholded comparisons. This yielded a structure-specific enrichment index:

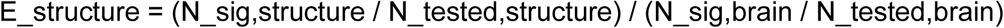

where N_sig,structure is the number of significant comparisons within a given neuropil, N_tested,structure is the total number of comparisons tested in that neuropil, N_sig,brain is the total number of significant comparisons across all neuropils, and N_tested,brain is the total number of comparisons across the complete dataset. Accordingly, E = 1 indicates that significant effects occur at the rate expected from the brain-wide background level, E > 1 indicates enrichment of significant modulation within that neuropil, and E < 1 indicates relative under-representation of modulation.

Because the medulla showed the strongest anatomical concentration of PSD modulation, we applied the same approach within this neuropil to determine which frequency bands contributed most strongly to the effect. For each frequency band, the fraction of significant medulla comparisons was normalized by the overall fraction of significant comparisons across all medulla bands:

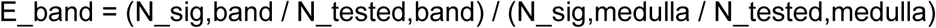

where N_sig,band is the number of significant medulla comparisons within a given frequency band, N_tested,band is the total number of medulla comparisons tested in that band, N_sig,medulla is the total number of significant comparisons across all medulla frequency bands, and N_tested,medulla is the total number of medulla comparisons across all bands. Here, E = 1 represents the expected medulla-wide level of modulation, E > 1 indicates over-representation of significant effects within that frequency band, and E < 1 indicates under-representation.

### Spike and MUA analysis

To assess how local neuronal output is related to luminance-dependent LFP dynamics in the medulla, we compared band-limited LFP activity with detected spikes and multi-unit activity (MUA) derived from the corresponding broadband extracellular recordings. Spikes were detected from the 300–5000 Hz band using a negative threshold of −4.5 SD relative to the median baseline and a 1.5-ms refractory period. Spike rate was subsequently calculated in 50-ms bins. MUA was extracted from the high-frequency signal by rectification followed by smoothing with a 10-ms moving average. MUA events were identified using a hysteresis-based detection procedure, with a threshold of +3.5 z to initiate an event and +2.5 z to terminate it, together with a minimum event duration of 5 ms and a 10-ms refractory period.

For LFP–spiking coupling analyses, LFP signals were band-pass filtered into the six data-driven frequency bands, and the amplitude envelope of each band was obtained using the Hilbert transform. Coupling between band-limited LFP amplitude and either spike rate or MUA was quantified using Spearman rank correlations, calculated both within individual trials and after pooling observations across trials. The squared Spearman correlation coefficient (ρ²) was used as a descriptive index of association strength between band-limited LFP activity and local spiking or MUA. Statistical comparisons between experimental conditions were performed using Welch’s t-tests for amplitude-based measures and Fisher z-transformed correlation coefficients for coupling measures, with false-discovery-rate (FDR) correction applied across frequency bands.

Analyses used custom MATLAB scripts. ChatGPT (OpenAI, GPT-5.4) assisted with writing, formatting, and code debugging. All outputs were manually verified by the authors.

### Histology

Brains were fixed in paraformaldehyde, washed in PBS, and cleared using a CUBIC-L (29) protocol at 30 °C for 4 wk. After DAPI staining and PBS washes, samples underwent ethanol-based dehydration and rehydration to reduce imaging artifacts. Brains were then treated with 50% and 100% RapiClear and stored at 4 °C. Imaging was performed with a Zeiss light-sheet microscope using 488-, 638-, and 405-nm laser lines. Volumes were stitched and visualized in Imaris to reconstruct probe trajectories and confirm neuropil assignments.

Because the probe followed a consistent lateral trajectory through the visual pathway and the geometry of the silicon probe was fixed and known, the anatomical identity of recording sites could be inferred from the combination of insertion path, contact spacing, and the ordered sequence of neuropils intersected by the electrode (retina, lamina, medulla, lobula, and protocerebrum). A representative histological reconstruction is shown for clarity and space considerations. This representative brain illustrates the same trajectory observed across preparations and confirms that the electrode crossed the expected visual neuropils in the predicted order. The known length and spacing of the recording contacts were then used to assign the analyzed sites along that trajectory.

### Data and code availability

Raw electrophysiological data generated in this study have been deposited in the Stockholm University Figshare repository under DOI 10.17045/sthlmuni.33436192. Behavioral data and the associated R analysis script are deposited separately under DOI 10.17045/sthlmuni.33461947. MATLAB code used for electrophysiological preprocessing and analysis is available under DOI 10.17045/sthlmuni.33102692

## Acknowledgments

This work was supported by the Swedish Research Council grants 2018-06238 and 2021-05564 (E.B.).

## Author Contributions

C.E.R.: Conceptualization; Investigation (electrophysiological experiments); Formal analysis (electrophysiological data); Software (electrophysiological analysis code); Writing – original draft and review & editing. P.A.: Investigation (behavioral experiments); Formal analysis (behavioral data); Writing – original draft and review & editing. S.M.: Software (electrophysiological analysis code). P.B.: Software (electrophysiological analysis code). E.B.: Conceptualization; Writing – original draft and review & editing.

---

Figure 1-figure supplement 1. Data-driven frequency bands identified from endogenous LFP activity. Mean global LFP power spectrum across all recordings, plotted as log10-transformed power as a function of frequency. Coloured segments indicate the frequency bins assigned to each cluster, and circles mark the mean frequency and mean power of each band identified via k-means clustering of log-normalized LFP power spectra.

Figure 2-figure supplement 1. Ocellar occlusion reorganizes luminance-dependent poststimulus activity relative to darkness. Heatmaps show differences in normalized local field potential power between each illuminated condition and the intact-dark reference across six recording sites and six data-driven frequency bands. Each panel compares the condition shown on the right with darkness on the left. Red indicates higher power in the illuminated condition, blue indicates lower power, and white indicates no thresholded difference. Saturated colors denote raw P < 0.05, and pale colors denote 0.05 ≤ P < 0.10; comparisons used Wilcoxon rank-sum tests. Counts are reported as [invariant, increased, decreased] region × band comparisons. (A) With the ocelli intact, dim-broadband stimulation produced a balanced mixture of increases and decreases across the visual pathway [12, 12, 12], demonstrating a distributed OFF-period response relative to darkness. (B) Bright-broadband stimulation shifted this pattern toward suppression [15, 7, 14], with decreases concentrated in the medulla and protocerebrum and additional effects in the lobula, whereas higher-frequency power increased in the retina. (C) Following ocellar occlusion, dim-broadband stimulation produced a more decrease-dominated pattern [11, 10, 15], particularly from 2.8 to 33.5 Hz in the medulla, lobula, and protocerebrum, together with increases at the lowest frequency and in the retina. (D) Under bright-broadband stimulation after ocellar occlusion, no thresholded differences from darkness were detected in the medulla across any frequency band. Residual modulation was concentrated mainly in the retina and in selected low- and mid-frequency bands at other recording sites [26, 4, 6]. Thus, the intact visual pathway generated distinct spectral responses to dim and bright stimulation, whereas ocellar occlusion reorganized these responses and shifted the medulla under bright stimulation toward a dark-like state. Re, retina; La, lamina; Me, medulla; Lo, lobula; ProL, lateral protocerebrum; ProM, medial protocerebrum

Figure 5-figure supplement 1. Frequency-resolved coupling between medulla multi-unit activity and local field potential under different luminance and ocellar conditions. (A) Representative medulla traces showing the high-frequency signal used for spike detection (top), the rectified and smoothed multi-unit activity (MUA) signal (middle), and the band-pass filtered LFP in the 33–45 Hz band together with its instantaneous amplitude envelope (bottom, orange). Black boxes indicate epochs enlarged in B. (B) Expanded examples of a threshold-detected spike event (top) and an MUA burst (bottom). (C) Strength of association between band-limited medulla LFP activity and MUA across six data-driven frequency bands under dark, dim-broadband, bright-broadband, dim-broadband-OC, and bright-broadband-OC conditions. Associations were quantified as squared Spearman correlation coefficients (ρ² × 100). Higher values indicate stronger covariation between MUA and the amplitude envelope of band-limited LFP activity.

