## Supplementary figures and images for "Dorsal ocelli set the luminance-dependent operating state of the bumblebee visual system"

### Supplemental Data 1

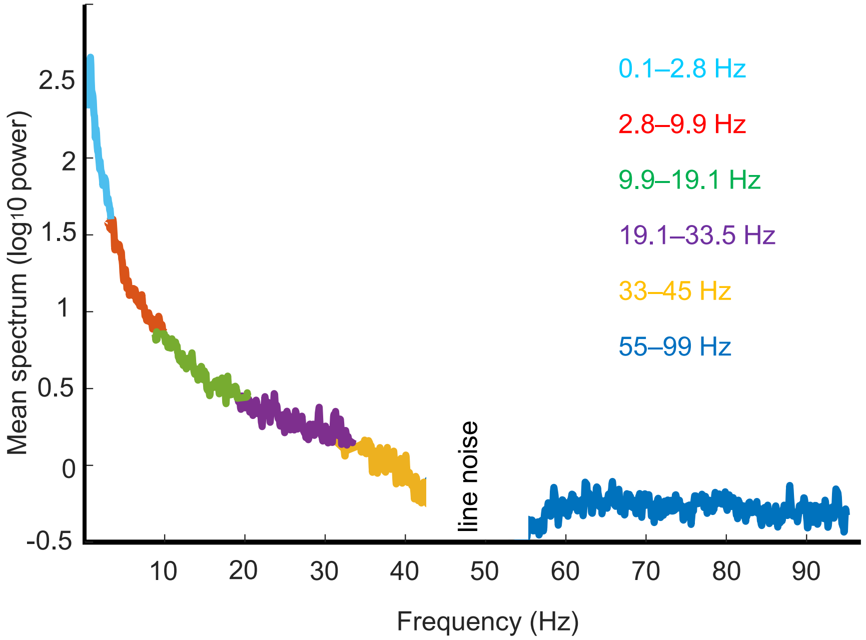

### Supplemental Data 2

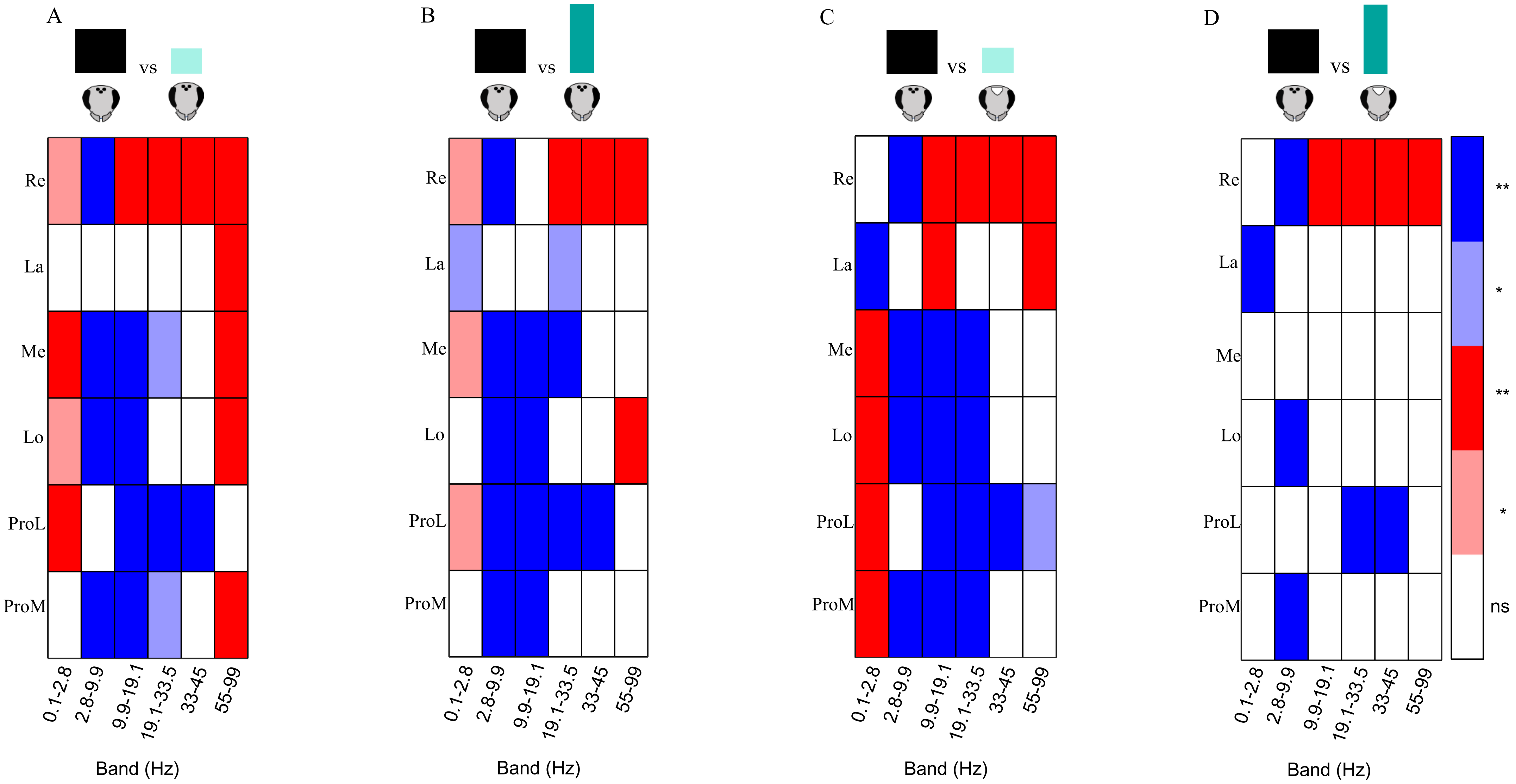

### Supplemental Data 3

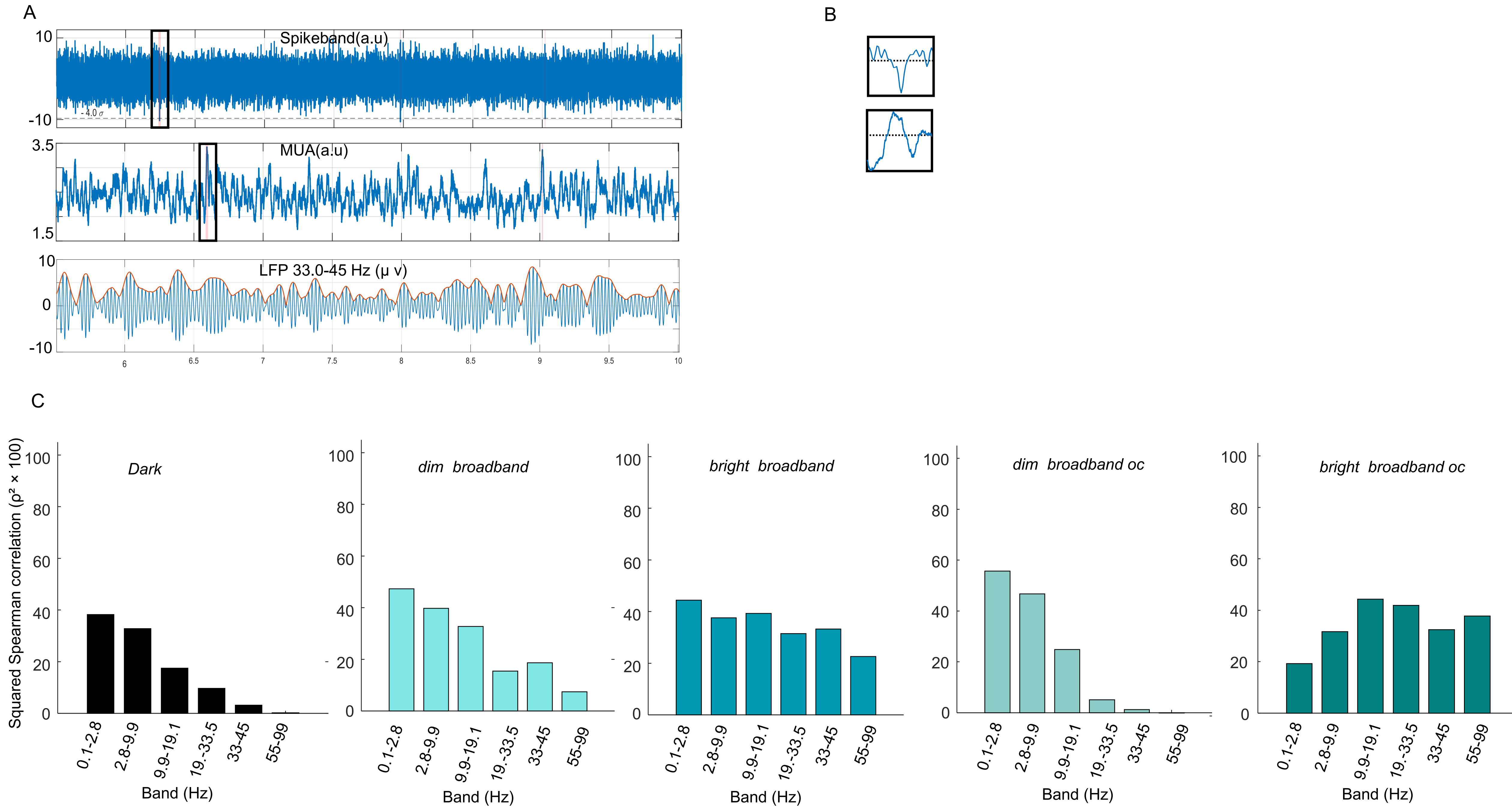
